# Leaf shape and size variation in bur oaks: An empirical study and simulation of sampling strategies

**DOI:** 10.1101/2020.05.11.088039

**Authors:** Sara C. Desmond, Mira Garner, Seamus Flannery, Alan T. Whittemore, Andrew L. Hipp

## Abstract

**PREMISE:** Leaf shape and size figure strongly in trees’ adaptation to their environments. Oaks are notoriously variable in leaf morphology. Our study examines the degree to which within-tree, among-tree, and among-site variation contribute to latitudinal variation in leaf shape and size of bur oak (*Quercus macrocarpa*: Fagaceae), one of North America’s most geographically widespread oak species.

**METHODS:** Samples were collected from four sites each at northern, central, and southern latitudes of the bur oak range. Ten leaf size traits were measured, and variance in these traits and eight ratios based on these traits was partitioned into tree, population, and latitude components. We then parameterized a series of leaf collection simulations using empirical covariance among leaves on trees and trees at sites. We used the simulations to assess the efficiency of different collecting strategies for estimating among-population differences in leaf shape and size.

**KEY RESULTS:** Leaf size measurements were highly responsive to latitude. Site contributed more than tree to total variation in leaf morphology. Simulations suggest that power to detect among-site variance in leaf shape and size can be estimated most efficiently with increases in either leaves per tree (10-11 leaves from each of 5 trees) or trees per site (5 leaves from each of 10+ trees).

**CONCLUSIONS:** Our study demonstrates the utility of simulating sampling and controlling for variance in sampling for leaf morphology, whether the questions being addressed are ecological, evolutionary, or taxonomic. Simulation code is provided as an R package (traitsPopSim) to help researchers plan morphological sampling strategies.

## INTRODUCTION

Leaf morphology variation strongly influences species’ ability to compete and survive in different environments (Givnish, 1987). The site-level correlation between temperature and the proportion of species exhibiting leaf dissection and toothing, for example, has been used for more than a century to model temperature changes in paleobotanical studies (Bailey and Sinnott, 1915, 1916; Greenwood et al., 2004; Royer and Wilf, 2006). Leaf size also has a well demonstrated correlation with temperature and resource availability (Bragg and Westoby, 2002; Peppe et al., 2011; McKee and Royer, 2017; Wright et al., 2017; Li et al., 2020), and traits such as compounding and phyllotaxy, leaf base and apex morphology, leaf lobing and circularity, and epidermal pigmentation vary along gradients of light availability, nutrient availability, soil moisture, temperature, and combinations of these (Givnish, 1987; Schmerler et al., 2012). In a global field study from 92 sites (Peppe et al., 2011), multiple regressions of climate (both precipitation and temperature) on leaf area, tooth number, and percent of species at the site with toothing showed relatively high predictive ability, inferred from the low standard error of the models (±4□). However, while the sign of this correlation—more toothing and lobing in cooler areas—is convergent across clades and geographic regions, the slope of the relationship between climate and leaf morphology varies among species (McKee et al., 2019), geographic regions (Greenwood et al., 2004; Aizen and Ezcurra, 2008), and phylogenetic lineages (Little et al., 2010; Burnham and Tonkovich, 2011; Walls, 2011).

Many of these traits vary both among and within species, and correlations between community-weighted mean trait values at the site level are mirrored within species on short time scales. Toothing and leaf lobing are, for example, correlated with cooler temperatures in most species studied as well as across communities (McKee and Royer, 2017; McKee et al., 2019). Moreover, the morphology of leaves can vary highly within communities (Givnish, 1987). Within forest trees in particular, leaves shape and size vary in response to leaf position on the tree (Blue and Jensen, 1988; McCarthy and Mason-Gamer, 2019), light availability (Abrams and Kubiske, 1990; Ducrey, 1992), and drought (Abrams, 1994; Abrams et al., 1994). Variation among leaves on a single tree is sometimes as great as the variation observed among named species (e.g., McCarthy and Mason-Gamer, 2019). Variation on a single tree is typically plastic, and much of the variation among trees at a site may be due to plasticity as well, but tree leaf morphology also varies in response to genetic differences among trees within species (Abrams, 1994; Ramírez-Valiente et al., 2017). While sampling of a small number of leaves per population has been argued to be sufficient for detecting site level patterns in climate based in paleobotanical studies (Royer et al., 2005; Peppe et al., 2011), the relative contribution of within-tree, among-tree within-population, and among-population variation to total leaf morphological variation is not known in many tree species.

Oaks exhibit highly variable morphology among leaves on a single tree, among trees of a single population, and among populations of a single species. Detailed studies of oak leaf morphology have utilized either linear measurements (Baranski, 1975; Blue and Jensen, 1988; Bruschi et al., 2003) or landmark approaches (Jensen, 1990). Both approaches have demonstrated that while variation among positions on a tree (e.g., high or low on the tree and disposed toward the edges or inside of the canopy) in leaf shape and size may exceed variation among sites, overall variance is generally greater among sites (Sokal et al., 1986; Blue and Jensen, 1988; Bruschi et al., 2003). Understanding these sources of variance in leaf shape and size is one key to understanding how introgression, adaptation, and neutral variation influence leaf morphology both among and within species (Jensen et al., 1984; Howard et al., 1997; Kremer et al., 2002; González-Rodríguez et al., 2004; González-Rodríguez and Oyama, 2005) and the balancing act that trees face in maximizing photosynthetic efficiency while minimizing the risks of drought, freezing, herbivory and other stresses (Wright et al., 2004). However, it is not clear what sampling strategy (number of leaves per individual, number of individuals per population, number of populations) is most efficient for estimating among-population differences in leaf size and shape. Whereas simulation tools exist for planning sampling strategies for population genetics (Hoban et al., 2013; Hoban, 2014) and conservation of genetic diversity (Hoban, 2019; Hoban et al., 2020), morphological simulation strategies that take into account covariance among leaves on a tree, among trees in a population, and among traits measured are lacking. Given that resources for sampling are limited, tools to help plan sampling strategies would make it possible to answer questions about functional plant diversity more definitively with the same amount of field work. The tools for planning morphometric sampling have not caught up with the tools for planning population genetic studies.

We sampled leaves across a broad geographic range of bur oak (*Quercus macrocarpa* L.), one of North America’s most geographically widespread oak species, which ranges from Manitoba to the Gulf of Mexico (Fig. 1). We measured leaf size and shape of numerous leaves per individual tree and per population to (1) quantify the relative contributions of within-tree, among-tree, and among-site variation to the total variation in leaf morphology in bur oak; (2) parameterize simulations of how much sampling is required to detect among-site differences in leaf morphology; and (3) test the support for our observations in field and herbarium that leaf size and leaf-lobing increase from north to south in bur oak. Bur oak serves as an excellent model species for this study because it has exceptionally high morphological variation (Hamerlynck and Knapp, 1994; Koenig et al., 2009) and an extensive distribution, ranging from Manitoba to the Gulf of Mexico (Little, 1971; Stein et al., 2003). The species also exhibits high within-population molecular genetic variation (Schnabel and Hamrick, 1990; Garner et al., 2019; Hipp et al., 2019). Our investigation of bur oak leaf morphological variation among vs within sites lays a foundation for our ongoing studies of what environmental factors contribute to functional variation in bur oak.

**Figure 1.**
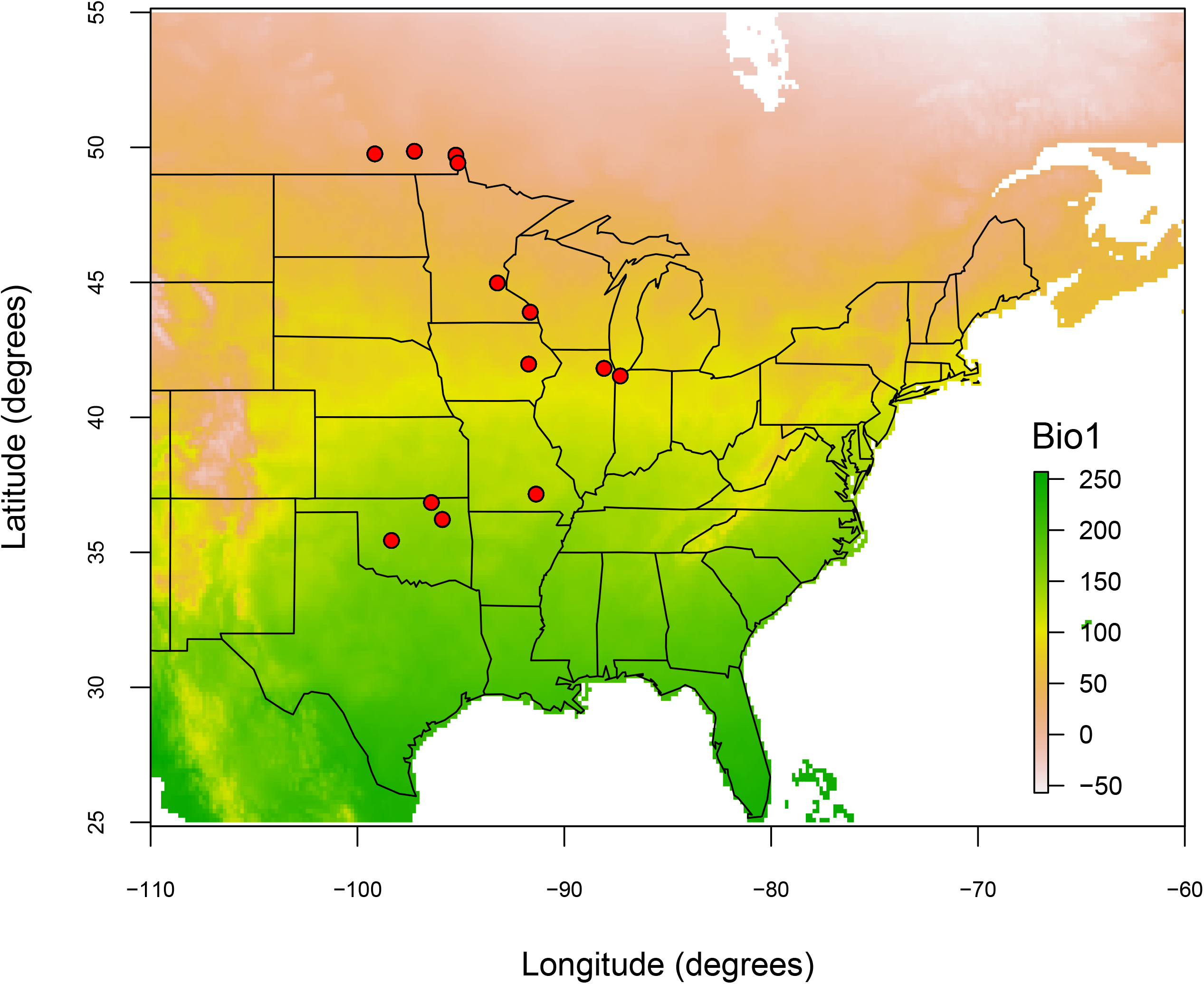
Locations of sampling sites for this study as well as mean annual temperature across the bur oak range. WorldClim temperature data are scaled to a factor of 10. Specific information about site, name, location, and number of samples collected can be found in Table 1.

## MATERIALS AND METHODS

### Collections and site attributes

During the summer and fall of 2017, samples were collected from four sites each at northern, central, and southern latitudes of the bur oak range (Fig. 1). Sites were selected in conjunction with sampling for population genetic studies of *Quercus macrocarpa* (Garner et al., 2019; Hipp et al., 2019) with the criteria that (1) preliminary collections suggested they would have numerous bur oaks, and (2) additional white oaks were present at the site or nearby. Most were forested, but some (e.g. The Morton Arboretum, Prairie Moon Nursery) were savannas. The northern sites sampled were located in Manitoba, Canada (Assiniboine Park, Whiteshell Provincial Park, and Spruce Woods Provincial Park) and Minnesota, U.S.A. (The University of Minnesota – Twin Cities). The central sites sampled were located in Illinois (The Morton Arboretum), Indiana (Burr Oak Woods), Iowa (Cherokee Park Trail), and Minnesota (Prairie Moon Nursery). The southern sites sampled were located in Oklahoma (Tallgrass Prairie Preserve, Mohawk Park, Red Rock Canyon State Park) and Missouri (Buttin Rock Access). Trees selected at each site were mature, full-size trees; where possible, trees at a given site were located a minimum of 100 feet from each other. For each site, latitude and longitude were recorded to a precision of 5 decimal places (Table 1). We extracted 19 bioclim variables from the WorldClim database (resolution = 1 km^2^) and linked them to our dataset in R v. 3.4.4 (R Core Team, 2018) using raster v 2.6-7 (Hijmans, 2017) and sp_1.4-2 (Pebesma and Bivand, 2005; Bivand et al., 2013) packages. The map of collection sites was made using maps 3.3.0 (Becker et al., 2018).

**Table 1.**
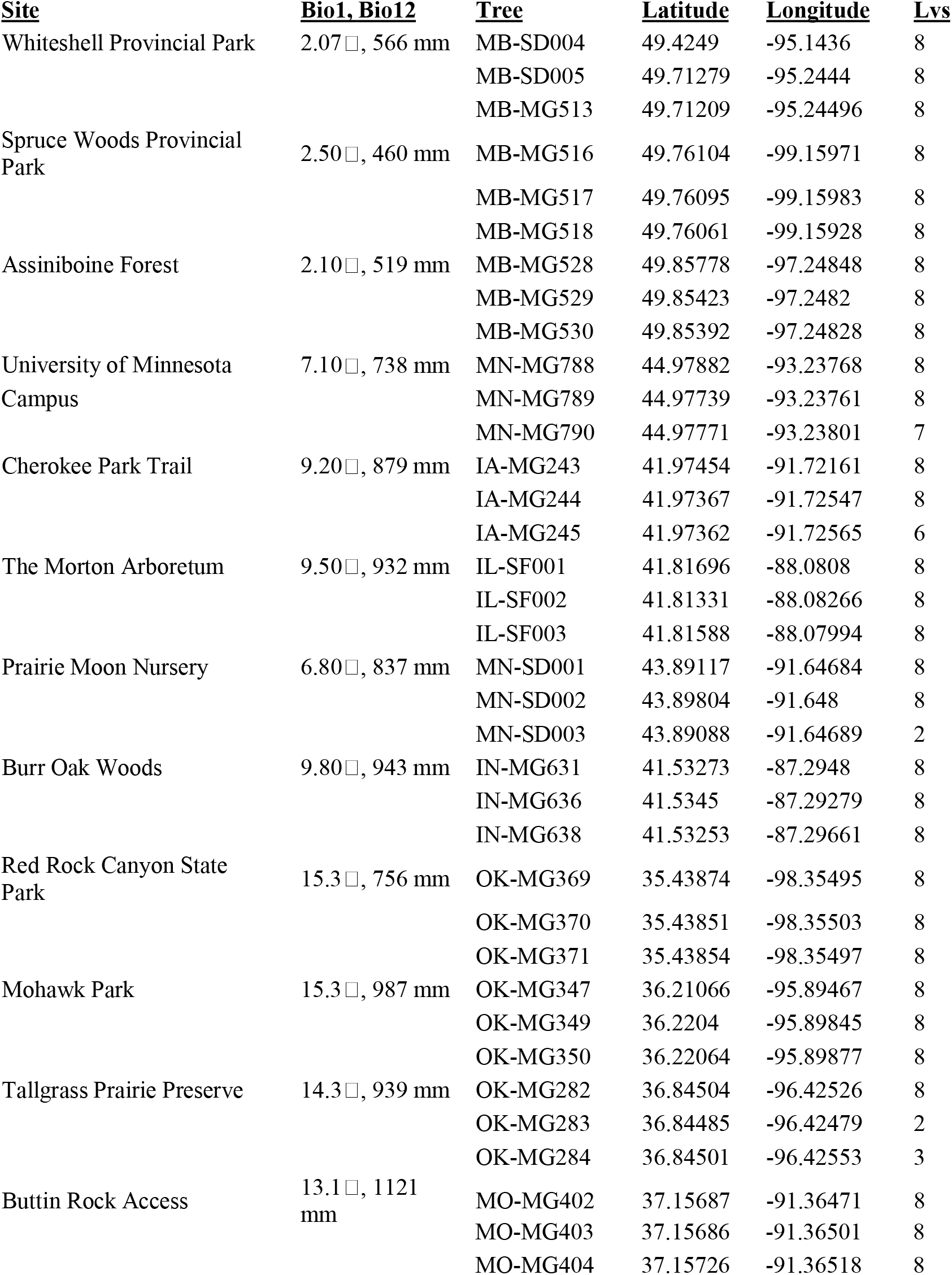
Sampling localities, Bioclim values for each site, number of leaves collected per tree. Only leaves used for statistical analysis are counted. Broken or incomplete leaves were eliminated from statistical analysis. Abbreviations: Bio1 = mean annual temperature (in degrees C); Bio12 = mean annual precipitation (in mm).

Three bur oak trees were sampled from each site using a pole pruner at two or four meters in height, based on tree height. For each sample, a terminal branch was cut down from each of the cardinal directions (N, S, E, W), determined using a compass. Only outermost branches were sampled. Two endmost leaves were removed from each branch and immediately pressed, for a total of 8 leaves per individual, 272 leaves overall. If the endmost leaves were highly damaged, the next leaves in from the end were selected. Leaves that were highly misshapen or broken were excluded from analyses (see ‘use’ field in dataset archived in GitHub repository [https://github.com/andrew-hipp/oak-morph-2020; https://doi.org/10.5281/zenodo.4458090], which indicates which leaves were excluded from analysis; all references to code and data in this paper are referable to the same GitHub repository). Leaves were dried in a standard herbarium drier prior to measuring, then redried at 49° C for a minimum of 48 hours and weighed on a PB303 Delta Range scale to obtain dry mass.

### Morphological Measurements

Ten size measurements (mm) were made on each leaf using ImageJ (Schneider et al., 2012): blade length (bladeL), blade width (bladeW), width of blade between deepest pair of sinuses (sinusMinW), petiole length (petioleL), petiole width (petioleW), length of lamina from base to widest point (bladeLtoWidestPoint), width of blade between pair of sinuses just above the deepest pair (sinusNextW), total length (BL.PL), leaf base angle (bladeBaseAngle), and leaf area (area) (Table 2, Fig. 2). Seven ratios were also calculated from these measurements to distinguish leaf shape from leaf size (González-Rodríguez and Oyama, 2005): petioleL / BL.PL (PL.TL); sinusMinL / sinusNextL (sinusRatio); bladeL / bladeW (BL.BW); petioleL / petioleW (PL.PW); BL.BW / PL.PW (BL.BW.over.PL.PW); bladeL / bladeLtoWidestPoint (BL.BLWP); lobedness, calculated as blade width between the deepest sinuses divided by total blade width, abbreviated (sinus.v.width); and specific leaf area (SLA), calculated as leaf blade area / leaf blade mass (Table 2). A panel of significant regressions was created using the R core functions and gridExtra 2.3 (Auguie, 2017).

**Table 2.** Descriptions of the leaf traits measured.

| <u>Leaf measurements</u> |  |  |
| --- | --- | --- |
| Trait | Abbreviation | Definition |
| Blade length (mm) | bladeL | Straight line distance measured from intersection of leaf and petiole to tip of the leaf at its point of intersection with the midvein |
| Blade width (mm) | bladeW | The longest possible perpendicular line drawn from one edge of the blade to the other; vein position used to identify the leaves opposite one another |
| Width of blade between deepest pair of sinuses (mm) | sinusMinL | The shortest distance that separates the deepest sinus from its corresponding sinus. (The deepest sinus is defined as the sinus that has the longest distance from the most interior point of the sinus to the line that connects the two most exterior points of that sinus) |
| Width of blade between sinuses just above the deepest pair (mm) | sinusNextL | The width between the sinuses that are immediately distal to the deepest sinuses (as defined in sinusMinL) |
| Petiole length (mm) | petioleL | Measured from the base of the blade as defined by bladeL to the base of the petiole, defined as the line of intersection between petiole and branch, upper surface of the petiole |
| Petiole width (mm) | petioleW | Measured at the point of intersection between the blade and the petiole, where blade is not visible |
| Length of lamina from base to widest point (mm) | bladeLtoWidestPoint | Measured from the base of the blade to the point of intersection between the midvein and the line used to measure leaf blade width |
| Blade base angle (degrees) | bladeBaseAngle | Measured using the lines that define the widest angle between the base and either edge of the leaf |
| Total length (mm) | BL.PL | Total length is the blade length added to the petiole length |
| Leaf area (mm<sup>2</sup>) | area | Calculated in imageJ. |

| <u>Ratios</u> |  |  |
| --- | --- | --- |
| Petiole length / Total length | PL.TL | The petiole length divided by the total length |
| Sinus ratio | sinusRatio | The width of the blade between the deepest pair of sinuses divided by the width of the blade between the pair of sinuses just above the deepest pair |
| Blade length/ Blade width | BL.BW | The blade length divided by the blade width |
| Petiole length / Petiole width | PL.PW | The petiole length divided by the petiole width |
| Blade length / Blade length to widest point | BL.BLWP | The blade length divided by the length of the blade from the base to the widest point |
| Ratio of Leaf size to petiole length | BL.BW.over.PL.PW | The blade length divided by the blade width divided by the petiole length divided by the petiole width |
| Specific leaf area | SLA | Leaf area divided by the mass of the leaf |
| Lobedness | sinus.v.width | The width of the blade between the deepest pair of sinuses divided by the blade width |

**Figure 2.**
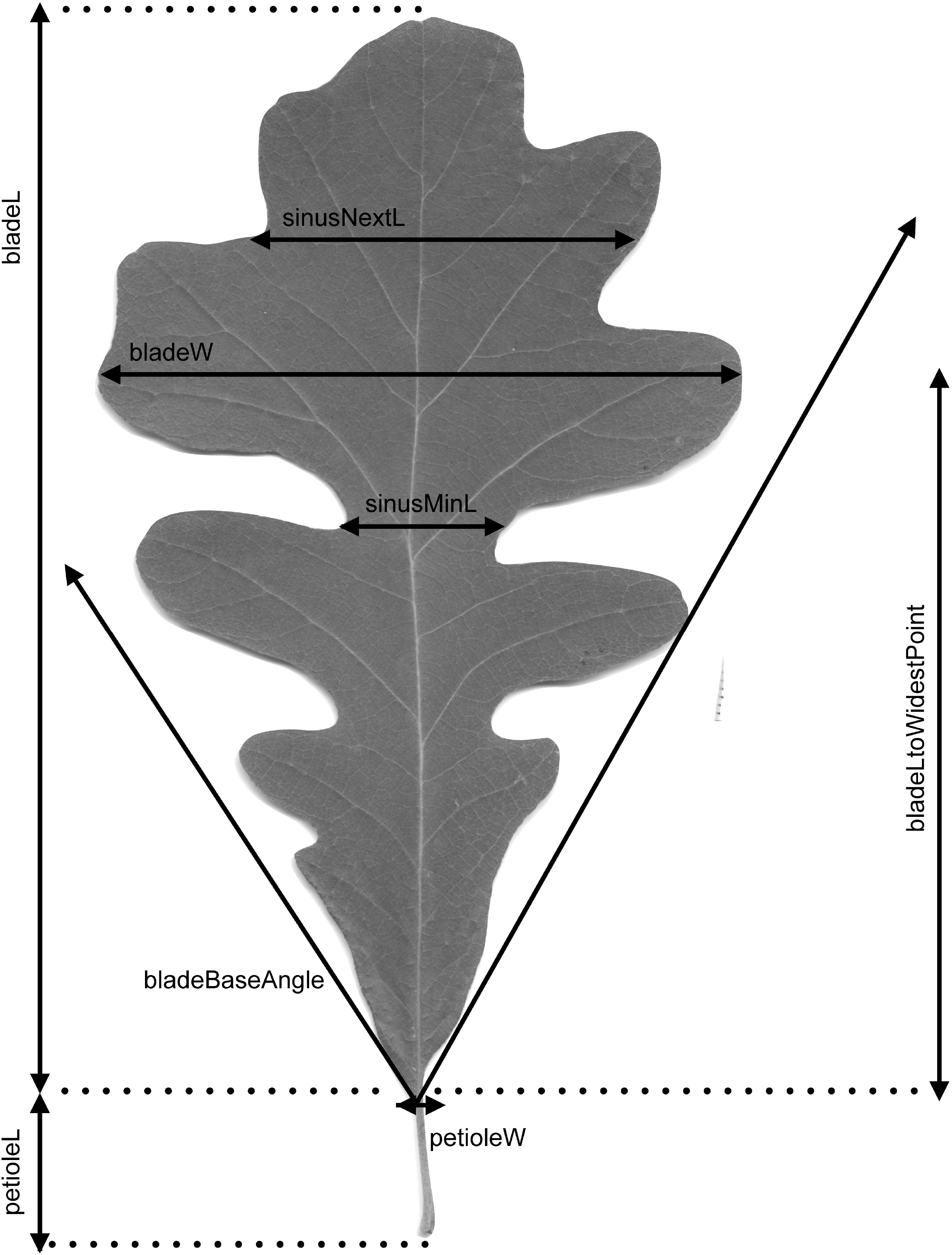
Leaf trait measurements used in this study. All measurements used in this study were linear measurements or ratios of linear measurements, plus one angle. Details and definitions are found in Table 4.

To investigate lobedness, black and white silhouettes of each leaf image were created using ImageJ and converted into jpeg files, with petioles whited out manually. The jpeg files were then imported into R and converted into outlines using the R package Momocs v 1.3.2 (Bonhomme et al., 2014). An additional set of 40 leaves that did not import well into Momocs or had leaf outlines that did not impair manual measurements but that were badly non-representative of typical leaf form were deleted at this stage (see scripts 05a and 05b in the GitHub repository for enumeration of these). We initially investigated shape variation using elliptical Fourier analysis (EFA), which generates shape-representative variables that are independent of size (Crampton, 1995) and is well suited to comparing complex outlines that vary in shape and lobedness (Tracey et al., 2006). For these analyses, we normalized the outlines using four landmarks placed on the top, bottom, left, and right of each outline and analyzed the leaf outlines using 17 harmonics (the default setting). However, even with landmarks, the leaf shapes were sufficiently complicated that the EFA did not yield useful results; they are consequently presented briefly in the paper; all data, code, and results are provided in the online supplement.

To further investigate lobedness and complexity of the leaf outline, we used Momocs to measure circularity (square of perimeter divided by area) (Rosin, 2005) and Haralick’s circularity (mean distance from the leaf centroid to perimeter pixels divided by standard deviation of those distances) (Haralick, 1974). Haralick’s circularity is less sensitive to shape raggedness than the standard measure of circularity and increases with increasing circularity; standard circularity decreases with increasing circularity. We aggregated both circularity measures to individual and then individuals to site to examine site-level effects of latitude on leaf circularity using simple least-squares regressions.

### Statistical Analysis

All statistical analyses were conducted in R. Least squares regressions were performed on all leaf linear measurements using the lm function to assess which leaf traits were most responsive to latitude at the site level, aggregating leaf traits first to tree, then tree mean trait values to site. An additional regression was performed of Haralick circularity on latitude, at the site level. Data were visualized using ggplot2 3.3.2 (Wickham, 2009). In addition to simple regressions, we corrected for size by conducting multiple regressions for all of our leaf traits using the lm function and including blade length (bladeL) as a covariate. We used data scaled to a mean of zero and unit variance so that regression coefficients estimate standardized effect sizes.

We performed a principal component analysis (PCAMORPH) on all scalar measurements and ratios using the prcomp function. The point MN-MG788 was removed prior to analysis because it significantly skewed the ordination. Two-dimensional nonmetric multidimensional scaling on a Euclidean distance matrix based on principal component axes was used to visualize the data. The scaling type was ‘centering’ with PC rotation. We used the ordiellipse function in vegan 2.4-5 (Oksanen et al., 2017) to plot bounding ellipses on our ordination. The resulting axes of the PCA are referred to hereafter in the paper as PC1_MORPH_ and PC2_MORPH_.

Two-way ANOVA was used to assess the relative contributions of site and tree to the total variation in bladeL, SLA, PC1_MORPH_ and PC2_MORPH_. ANOVA was conducted on the linear model of bladeL, SLA, PC1_MORPH_, and PC2_MORPH_ regressed against site and tree. We chose PC1_MORPH_ and PC2_MORPH_ for this analysis because together they accounted for 52.5% of the variance.

### Simulations of sampling strategies

We assessed the effectiveness of alternative sampling scenarios at distinguishing differences in leaf shape and size among populations by using our empirical data to parameterize simulations of morphological datasets collected from 20 populations, ranging from three to 12 trees per site and three to 12 leaves per tree, a total of 100 sampling strategies. For each strategy, we simulated 100 replicate datasets of all ten direct morphological measurements using a hierarchical simulation strategy. For each replicate, site-level means for all 10 traits were drawn from the multivariate normal distribution with trait means and covariance **C**_site_ estimated from empirical site means for all traits; **C**_site_ is thus based on variance within and covariance among traits that we observed, averaged for each site. Tree-level means were then drawn from the multivariate normal distribution with the simulated site-level means and the covariance matrix **C**_tree_ estimated from tree means at each site and averaged across sites. Tree-level means were assumed to have a constant variance and covariance among sites, though this assumption could be relaxed. Finally, individual leaf measurements for each tree were drawn from the multivariate normal distribution with means from the second simulation stage and covariance matrix **C**_leaf_ estimated from the leaf measurements for each tree separately, then averaged across trees.

The simulated data matrices ranged from 180 to 2,880 simulated leaves, with trait covariance and variance among leaves within trees, among trees within populations, and among populations modeled according the measurements we made for this project. Because leaf size showed particularly strong variation among populations, we utilized ANOVA of bladeL on site + tree, combined with Tukey’s Honest Significant Different (HSD) method to assess the number of populations that could be differentiated from one another in each simulated data matrix. The number of letters needed for a compact letter display using Tukey’s HSD at α = 0.05 was used as a proxy for the number of groups that could be distinguished for each simulated dataset. Both the average number of groups distinguished for each simulated dataset and the percent of simulations that distinguish at least 50% of populations (at least 10 groups distinguished from the 20 populations simulated) are reported as estimates of statistical power. All simulations were conducted in R using mvtnorm v 1.0-11 (Genz and Bretz, 2009; Genz et al., 2019) (Genz et al. 2017), and code for performing simulations is archived in GitHub and Zenodo (https://github.com/andrew-hipp/oak-morph-2020; https://doi.org/10.5281/zenodo.4458090). The simulation code can be installed as package ‘traitsPopSim’ from GitHub (https://github.com/andrew-hipp/traitsPopSim) and run using multivariate traits collected in a similarly structured design (measurements nested within individuals nested within sites; sample data included in the package are from the current study).

## RESULTS

### Analysis of empirical data

Among the size characters, bladeL, bladeW, BL.PL, petioleW, and area all showed significant variation in response to latitude (Table 3, Fig. 3). Petiole width was the size trait that was the most significant (P < 0.001). Petiole length, sinusMinL, sinusNextL, and blade base angle were not significantly predicted by latitude (Table 3). Two ratios were significantly correlated with latitude: SLA (*r*^2^ = 0.466, P = 0.015) and the ratio of sinus depth to leaf width (sinus.v.width, a proxy for lobedness; *r*^2^ = 0.415, P = 0.024) (Table 3). In regressions that include bladeL as a covariate, sinusNextL (P = 0.021), petioleW (P = 0.017), and SLA (P = 0.011) were the only traits significantly affected by latitude (Table 3; Fig. 3). Biplots with fitted lines for all traits measured, significant or not, are presented in the supplement (Appendix S1; see the Supplementary Data with this article). Standard circularity (perimeter squared over area) was strongly correlated with latitude (*r*^2^ = 0.593, P = 0.0034; Fig. 4). Haralick circularity (mean distance from leaf centroid to boundary pixels over the standard deviation of these distances) was weakly correlated with latitude (*r*2 = 0.256, P = 0.0936; Fig. 4). Under both measures, southern leaves exhibited stronger lobing, northern leaves exhibited greater circularity.

**Table 3.**
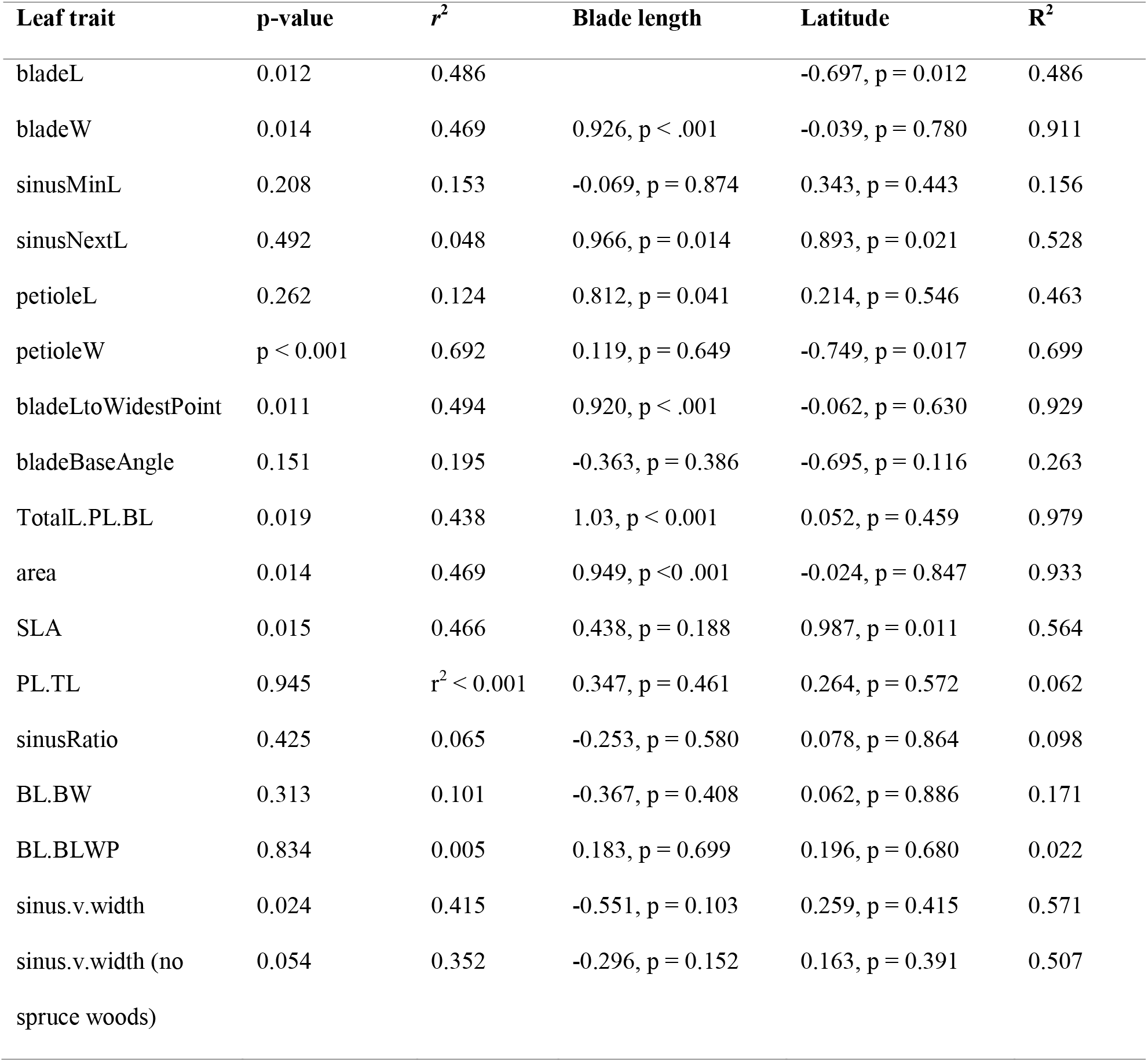
Simple and multiple regressions for all leaf traits. The p-value and *r*^2^ columns indicate the simple regression of each trait on latitude; the columns labelled Blade length and Latitude represent the regression coefficient and p-value for a multiple regression with each leaf trait regressed against Blade length and Latitude, *R*^2^ indicates the multiple coefficient of determination. Note that Bonferroni correction for multiple tests, not shown here, would render all p-values non-significant except the regression of blade length on latitude + petiole width.

| Leaf trait | p-value | $r^2$ | Blade length | Latitude | $R^2$ |
| --- | --- | --- | --- | --- | --- |
| bladeL | 0.012 | 0.486 |  | -0.697, p = 0.012 | 0.486 |
| bladeW | 0.014 | 0.469 | 0.926, p < .001 | -0.039, p = 0.780 | 0.911 |
| sinusMinL | 0.208 | 0.153 | -0.069, p = 0.874 | 0.343, p = 0.443 | 0.156 |
| sinusNextL | 0.492 | 0.048 | 0.966, p = 0.014 | 0.893, p = 0.021 | 0.528 |
| petioleL | 0.262 | 0.124 | 0.812, p = 0.041 | 0.214, p = 0.546 | 0.463 |
| petioleW | p < 0.001 | 0.692 | 0.119, p = 0.649 | -0.749, p = 0.017 | 0.699 |
| bladeLtoWidestPoint | 0.011 | 0.494 | 0.920, p < .001 | -0.062, p = 0.630 | 0.929 |
| bladeBaseAngle | 0.151 | 0.195 | -0.363, p = 0.386 | -0.695, p = 0.116 | 0.263 |
| TotalL.PL.BL | 0.019 | 0.438 | 1.03, p < 0.001 | 0.052, p = 0.459 | 0.979 |
| area | 0.014 | 0.469 | 0.949, p < 0.001 | -0.024, p = 0.847 | 0.933 |
| SLA | 0.015 | 0.466 | 0.438, p = 0.188 | 0.987, p = 0.011 | 0.564 |
| PL.TL | 0.945 | $r^2 < 0.001$ | 0.347, p = 0.461 | 0.264, p = 0.572 | 0.062 |
| sinusRatio | 0.425 | 0.065 | -0.253, p = 0.580 | 0.078, p = 0.864 | 0.098 |
| BL.BW | 0.313 | 0.101 | -0.367, p = 0.408 | 0.062, p = 0.886 | 0.171 |
| BL.BLWP | 0.834 | 0.005 | 0.183, p = 0.699 | 0.196, p = 0.680 | 0.022 |
| sinus.v.width | 0.024 | 0.415 | -0.551, p = 0.103 | 0.259, p = 0.415 | 0.571 |
| sinus.v.width (no spruce woods) | 0.054 | 0.352 | -0.296, p = 0.152 | 0.163, p = 0.391 | 0.507 |

**Figure 3.**
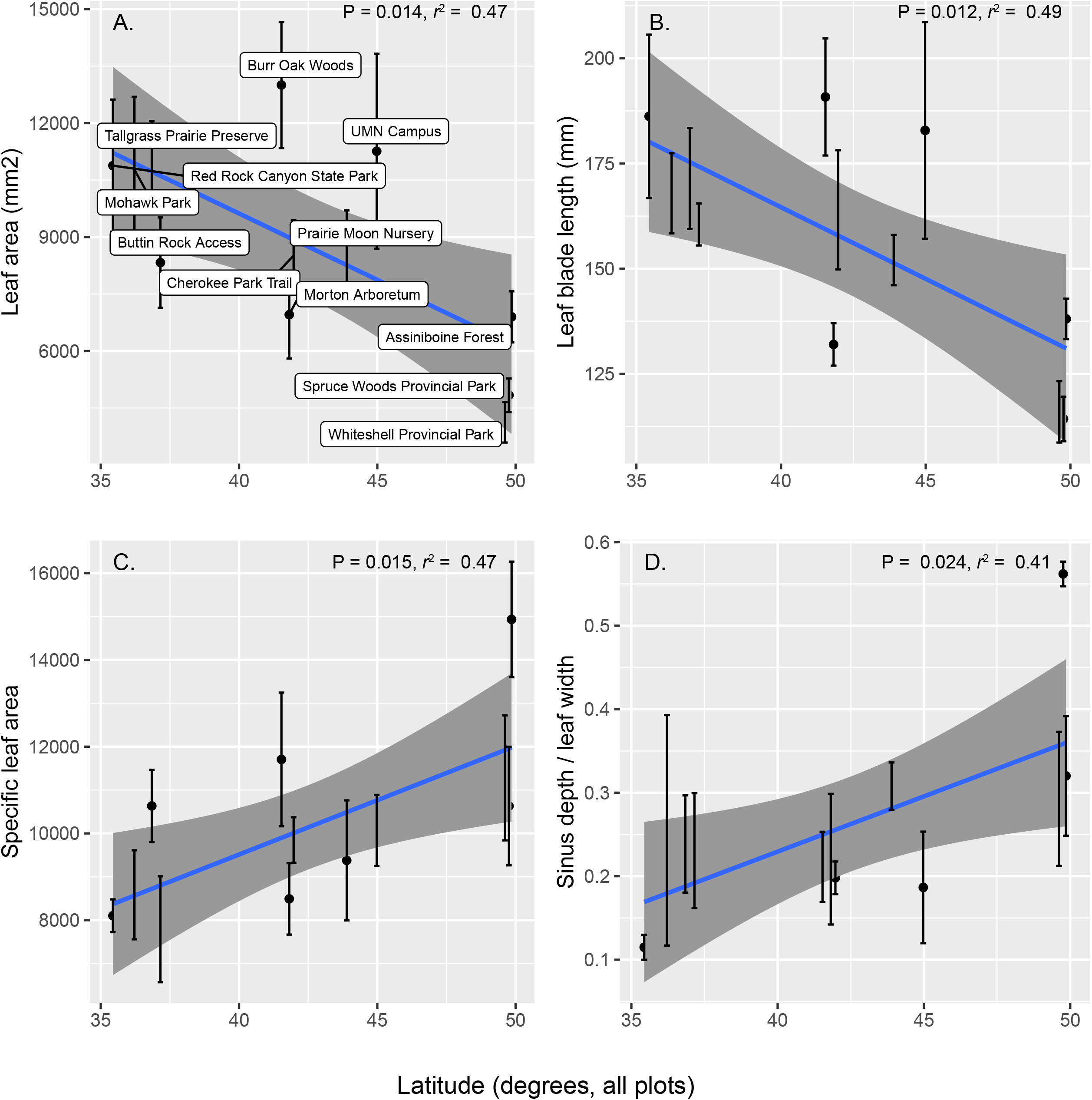
Simple regressions of traits and environment that are significant at the 0.05 level. P-values are not corrected for multiple tests; a total of seventeen regressions were performed (Table 3).

**Figure 4.**
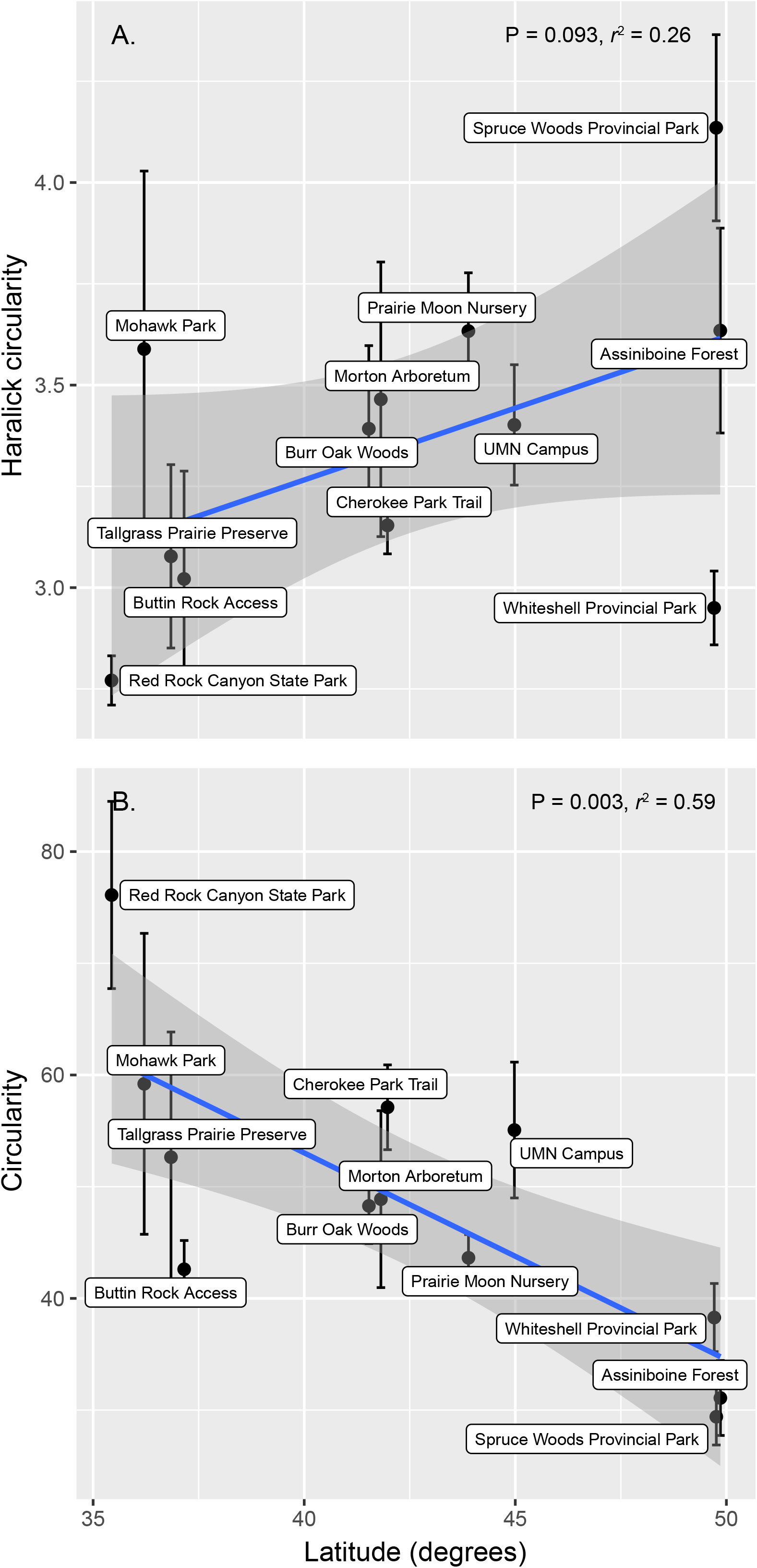
Simple regressions of circularity on latitude.

PCA on the EFA yielded very high loading on PC1 (58%), very low on PC2 (8%), and a strong curvilinear relationship between PC1 and PC2 (Appendix S2). This non-independence between PC1 and PC2 can arise because variation is broad, such that the ends of the PC cloud have little in common, or because the variation is dominated by a single variable (Minchin, 1987). In our study, PC1 was dominated by size (Pearson’s product moment [*r*] = 0.251 for blade width, 0.227 for blade area) and latitude (*r* = −0.213), and PC2 was more strongly associated with shape (*r* = 0.395 for the ratio of blade length to blade width, −0.375 for width of the sinus distal to the deepest sinus, 0.348 for the ratio of the width of the blade in the deepest sinus width of the blade in the second-deepest sinus), and to a lesser extent leaf blade width (*r* = −0.240).

The effects of site and tree on bladeL, SLA, and the first two axes of the morphological (non-EFA) ordination (Appendix S3) were significant based on ANOVA (P << 0.001; Table 4). Although site and tree both had significant effects, site contributed more than tree to the total variation in leaf morphology (F-values for site range from 30.38-41.76, while F-values for tree range from 5.83-12.4). Mean annual temperature among our sites ranged from 2.1–15.3°C, and mean annual precipitation from 460 – 1121 mm. On average, bladeL averaged 34.0 mm shorter and SLA 50.39 mm^2^/g greater for each increase of 10 degrees in latitude (northward).

**Table 4.**
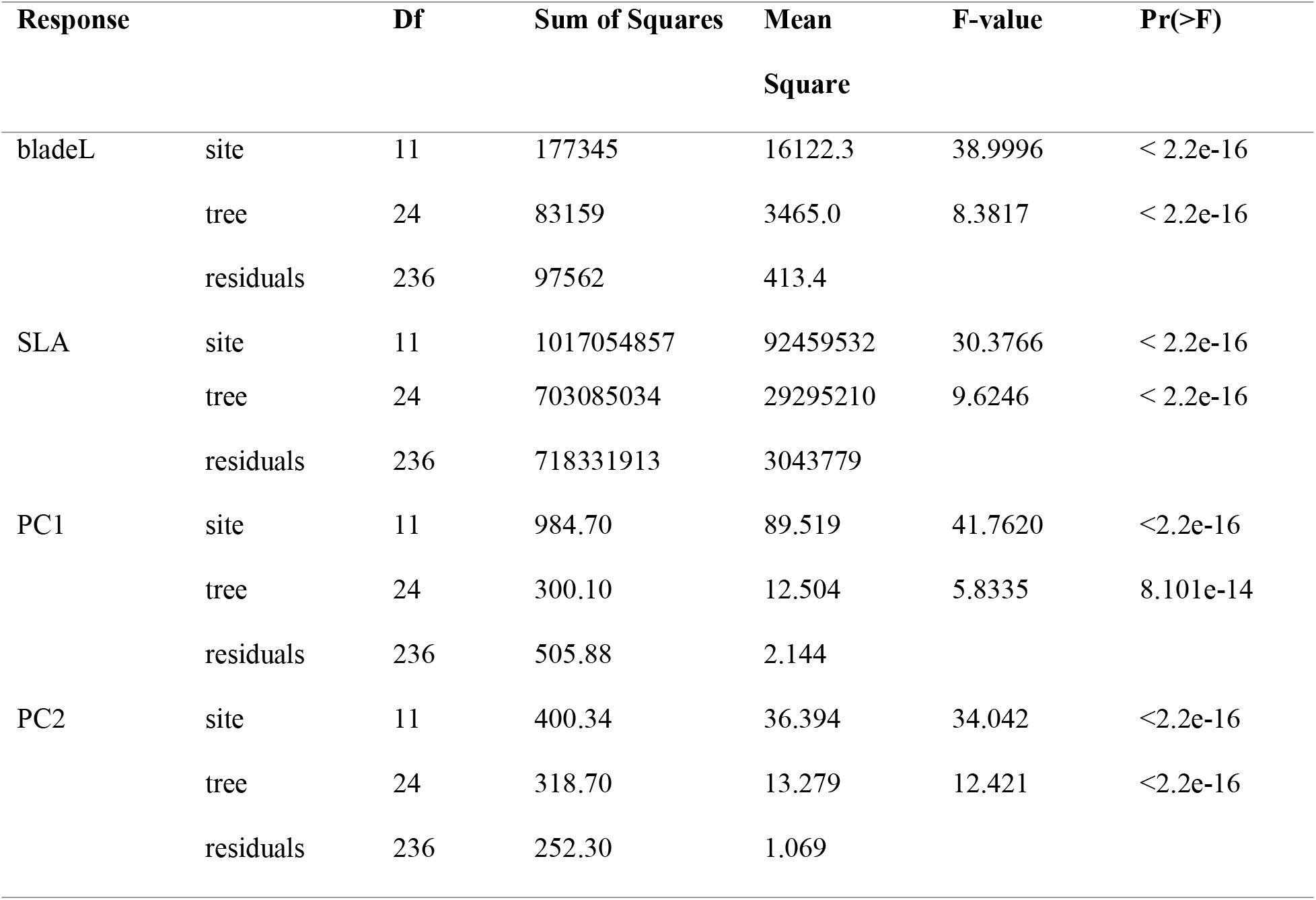
ANOVA for bladeL, SLA, PC1, PC2.

Regressions of individual traits on site-level temperature and moisture conditions inferred from BioClim closely matched regressions of those same traits on latitude. The latitudinal gradient in our study correlated tightly with climate: increases in latitude correlated with decreases in mean annual precipitation (Bio12; *R*^2^ = 0.6803, p < 0.01) and temperature (Bio1; *r*^2^= 0.99, p << 0.01), and an increase in temperature seasonality (Bio4; *R*^2^ = 0.97, p << 0.01) (Fig. 5). As a consequence, climate is not considered further in this study, but only latitude.

**Figure 5.**
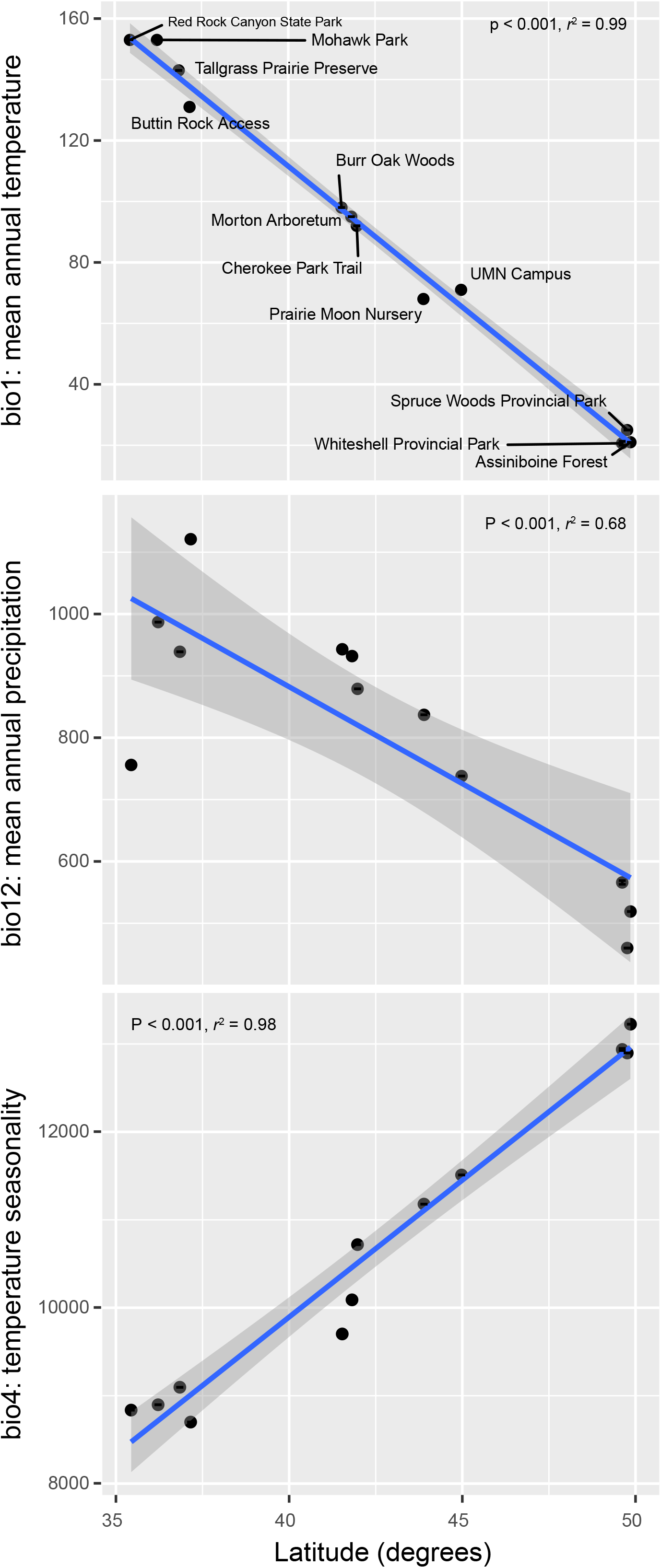
Regressions of bioclim variables on latitude. Latitude shows up as the strongest single predictor of leaf morphology in the current study, as it integrates over both day length and several aspects of climate: bio 1 (mean annual temperature), bio 12 (mean annual precipitation), bio 4 (mean temperature seasonality).

### Analysis of simulated data

The mean number of groups distinguished in our simulations ranged from 5.49 to 10.41, and the probability of distinguishing 50% (10 / 20) of the populations based on blade length ranged from 0.01 to 0.71 (Fig. 6). The sampling strategy we implemented for this study, 3 trees per site, 8 leaves per tree, had a power of only 38% to identify a number of groups equal to 50% of the sites sampled. Increasing power to at least 50% would entail increasing sampling to 11–12 leaves from each of 5 trees, 5 leaves from each of 10–11 trees, or any of a number of scenarios intermediate between these extremes.

**Figure 6.**
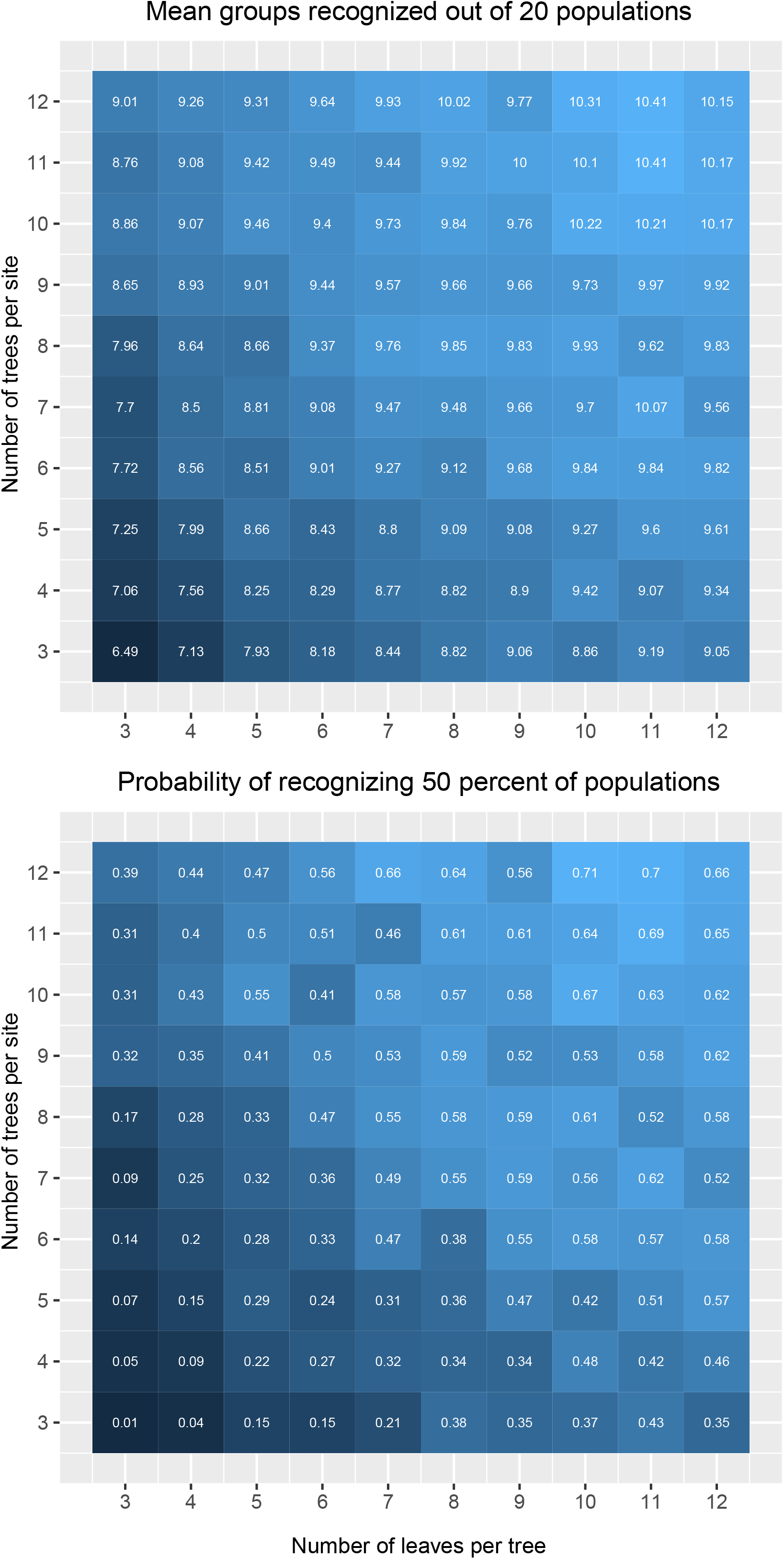
Sampling simulations. Simulated sampling strategies accounted for covariance among traits within leaves; among leaves on trees within sites; and among trees within sites. Here, two estimates of power are reported: the number of groups of sites recognized as distinct from each other using Tukey’s HSD at α = 0.05; and the probability of recognizing at least 50% of sites as distinct from each other. colors scale from darker as a higher number of groups are recognized, lighter as fewer are. Simulated numbers of sites distinguished (left panel) and probabilities of distinguishing at least 50% of simulated sites (right panel) are reported in each cell of the simulation.

## DISCUSSION

Our study has three important findings. First, among-tree and among-site variation contribute significantly to leaf shape and size variation in bur oaks. Consequently, within-individual and within-population sampling are both important components of a sampling strategy aimed at characterizing among-population variation in oak morphology. This complements observations of high variance in temperate tree leaf morphology (Bruschi et al., 2003; Apostol et al., 2017; McCarthy and Mason-Gamer, 2019): among-site variation generally contributes most strongly to total leaf variation, but within-site and within-tree sampling are important to detecting among-site variation in leaf shape and size. Second, we implement a general parametric simulation method and use it to demonstrate that our sampling strategy, which included 8 leaves from different positions on each of 3 trees per site, was not optimal for resolving among-site variation, even if it was sufficient to demonstrate the relationship between morphology and latitudinal gradients. This simulation approach and the R package provided (traitsPopSim; https://github.com/andrew-hipp/traitsPopSim) can serve as tools to guide morphological sampling in similar hierarchical studies, where sites are composed of multiple individuals and individuals are each represented by multiple measurements. Finally, our study demonstrates that leaf size and lobing decrease from south to north in bur oak, while specific leaf area increases. In cross-species comparisons, leaf size and SLA generally covary, suggesting that adaptive leaf variation in bur oak may rest in part on a tradeoff between leaf size and lobing. Our study of one of North America’s most widespread oak species is thus a jumping-off point for understanding adaptive leaf variation across the oaks of the Americas.

### Leaf size and shape are influenced by population and individual

Our results show that among-site variance for all traits investigated (F_11,236_ = 30.38–41.76) contributes more to total variance in leaf morphology than among-tree variance (F_11,236_ = 5.83–12.42), though both variance components are significant (P << 0.001; Table 4). This ability to distinguish among sites is important in relating leaf variation to latitude or climatic predictors and measuring the slope of the relationship resulting from selective pressures along climatic gradients (Wright et al., 2004). Moreover, our results demonstrate that sampling three trees per site and eight leaves per tree is sufficient to correlate shape and size to latitude and climate. However, while among-site variance is higher than among-tree variance within sites (Table 4), the variance we observe among leaves within a single tree is still quite high. A previous study (Bruschi et al., 2003) found that among-leaf morphological variance on a tree was higher than among-tree variance for most traits investigated and reported that this was in accord with findings from earlier work (Baranski, 1975; Blue and Jensen, 1988). However, in Bruschi (2003), leaves were sampled from both inner and outer positions on the branch to maximize variance. In our study, we deliberately minimized this source of variance by sampling leaves at a relatively constant height and all from the outer branch position, and we further selected the endmost leaves from each branch sampled. Unless researchers are interested in characterizing variation among leaves on single trees, we recommend minimizing within-tree variance by standardizing the position(s) on the tree from which leaves are collected.

### Simulating sampling strategies

The simulations we conducted of alternative sampling strategies suggest that the strategy we selected of three trees per site and eight leaves per tree has only a 38% probability of distinguishing 50% of the 20 populations we simulated. This is based on variances and covariances among trees within sites, leaves within trees, and covariances among the traits we measured, estimated from our empirical data. Based on the variance observed in leaf length alone, achieving a 50% probability of distinguishing 50% of populations would require 11–12 leaves from each of 5 trees per site, 5–6 leaves from each of 11–12 trees per site, or something in between (Fig. 6).

Our simulations suggest two recommendations for others conducting similar studies. First, researchers are recommended to minimize the high within-individual variance observed in previous studies (Blue and Jensen, 1988; Bruschi et al., 2003; McCarthy and Mason-Gamer, 2019) by sampling leaves of a common age / developmental stage, in the same position on the twig, and from twigs with comparable positions on the plants. Second, simulating alternative sampling strategies will help maximize the ability to distinguish among populations, given limited time and resources. Researchers can use preliminary data to simulate alternative sampling strategies and estimate their power will be to distinguish populations under different scenarios. The simulation tool implemented in traitsPopSim requires only a matrix of traits and assignment of those traits to populations and individuals to perform the simulations we describe above. We hope that its use will facilitate planning of sampling designs for similar projects.

### Leaf size, lobing, and SLA are predicted by latitude

The measurements for each of our leaves were well predicted by their latitude of origin: leaves were thicker, larger, and had deeper lobes at southern latitudes, where leaves are exposed to warmer temperatures and higher precipitation and have longer growing seasons; and leaves were smaller, thinner, and had shallower lobes at northern latitudes, where growing seasons are shorter and cool temperatures reduce drought stress. This is in line with previous studies demonstrating that leaf area covaries positively with temperature (Moles et al., 2014; Wright et al., 2017) while SLA covaries negatively (Moles et al., 2014), and that leaf circularity tends to increase in more northern or cooler environments (Halloy and Mark, 1996; Schmerler et al., 2012). Our results also parallel previous work in *Quercus ilex*, which exhibited a similar leaf size gradient from north to south in the western Mediterranean basin, where southern regions were likewise warmer and had higher amounts of precipitation than northern regions (García□Nogales et al., 2016).

Our findings suggest a possible compensatory relationship between larger size and lobing in bur oak. Community-level studies tend to show a higher frequency of lobed leaves in cooler temperatures (Royer et al., 2005). These responses are individualistic, however, and among-population responses in some species show no response or greater lobing in warmer temperatures (Royer et al., 2008; McKee and Royer, 2017; McKee et al., 2019). Leaves that are deeply lobed may be better adapted to warmer climates, because deeply lobed and narrow leaves have a thinner leaf boundary layer, facilitating more rapid cooling (Givnish, 1987; McDonald et al., 2003). In our study, the ratio of sinus depth to leaf width (sinus.v.width) shows a weak negative correlation with latitude (*b* = 0.013, P = 0.024), but only in simple regression. Moreover, this result is strongly affected by one site, Red Rock Canyon, which had an exceptionally low value. When this outlier is removed, the correlation is no longer significant (*b* = 0.007, P = 0.054). In multiple regressions with scaled data and bladeL as a covariate, however, depth of the sinus immediately above the deepest sinus (sinusNextL) was significantly influenced by latitude (*b* = 0.893, P = 0.021), even with the outlier removed (*b* = 0.926, P = 0.023). Our whole-leaf estimates of shape (circularity and Haralick circularity) similarly both showed increased circularity northward, but their sensitivity to the latitudinal gradient is different. This may be a consequence to the relative insensitivity of Haralick circularity to leaf toothing (Haralick, 1974), which manifests in bur oak as differences in crenulation. We did not quantify this effect directly, but leave it to future studies.

The hypothesis that leaf lobing increases southward in response to increased drought stress is supported by the specific leaf area (SLA) data. With leaf length as a covariate, SLA increases northward (*b* = 0.987, P = 0.011), even as leaf size decreases. Leaves that are low in SLA have higher water use efficiency (Mooney and Dunn, 1970; Marron et al., 2003; Liu et al., 2017), and SLA has been shown to vary within oaks according to drought stress (Ramírez-Valiente and Cavender-Bares, 2017; Ramírez-Valiente et al., 2017). However, leaf area in cross-species oak comparisons covaries with SLA (Ramírez□Valiente et al., 2020), and leaf area and SLA both decrease on average with increased water stress in cross-species comparisons (Kaproth and Cavender-Bares, 2016; Ramirez Valiente et al., 2020). Our finding of larger leaf areas with lower SLA in bur oak, but with an increase in lobing, shows the opposite correlation. We hypothesize that among populations within bur oak, leaf lobing compensates for increased size as a response to drought stress and temperature.

The immense success of oaks (*Quercus*) in the Americas (Rodríguez-Correa et al., 2015; Hipp et al., 2018; Cavender-Bares, 2019) has been attributed in part to oaks’ ability to cross the temperate-tropical divide. Bur oak is exceptional in its climatic range, extending from near the boreal zone in the north to the great plains and the humid subtropics. Our inference that leaf lobing, SLA, and leaf size may compensate for one another along climatic gradients in bur oak may be echoed in other species. The work presented here consequently has the potential to inform studies of adaptive variation across oaks and temperate tree species more generally.

## ACKNOWLEDGMENTS

This study was funded by The Morton Arboretum Center for Tree Science and USDA Project 8020-21000-070-03S, a non-assistance cooperative agreement between U.S. National Arboretum and The Morton Arboretum. We would like to thank Marlene Hahn for assisting with curation of herbarium and leaf samples and Matthew Kaproth for advice on measurement of SLA. Ricardo Kriebel provided particularly detailed advice on morphometric analysis, including code to assist in generating landmarks and excellent feedback on interpretation of our results and Associate Editor Dylan Schwilk, with an anonymous reviewer and Kriebel, provided editorial feedback that substantially improved the manuscript.

## AUTHOR CONTRIBUTIONS

A.L.H., A.T.W. and S.C.D. conceptualized and designed the project. S.C.D., M.G., and S.F. collected specimens and data. S.C.D. and A.L.H. conducted data analyses. S.C.D. wrote the first draft of the manuscript, and A.L.H. revised the manuscript in response to reviewers. A.L.H. coded and analyzed simulations and contributed to data analysis. All authors contributed to writing and revisions; A.L.H. and S.C.D. contributed equally to analysis and writing.

## DATA ACCESSIBILITY

Data and scripts used to conduct the statistical analysis are archived in GitHub (https://github.com/andrew-hipp/oak-morph-2020) and released through Zenodo (https://doi.org/10.5281/zenodo.4458090). Simulations are packaged for R in the traitsPopSim package, published on GitHub (code and installation instructions at https://github.com/andrew-hipp/traitsPopSim) and released through Zenodo (https://doi.org/10.5281/zenodo.4457926). Supplemental figures are available in the GitHub repository and in the online Supplement.

## SUPPORTING INFORMATION

Additional Supporting Information may be found online in the supporting information section at the end of the article.

Appendix S1. Biplots of all simple regressions

Appendix S2. PCA based on eFourier analysis of leaf outlines

Appendix S3. Non-metric multidimensional scaling ordination of leaf measurements

## REFERENCES CITED

Abrams, M. D. 1994. Genotypic and phenotypic variation as stress adaptations in temperate tree species: a review of several case studies. Tree Physiology 14: 833–842.

Abrams, M. D., and M. E. Kubiske. 1990. Leaf structural characteristics of 31 hardwood and conifer tree species in central Wisconsin: Influence of light regime and shade-tolerance rank. Forest Ecology and Management 31: 245–253.

Abrams, M. D., M. E. Kubiske, and S. A. Mostoller. 1994. Relating Wet and Dry Year Ecophysiology to Leaf Structure in Contrasting Temperate Tree Species. Ecology 75: 123–133.

Aizen, M. A., and C. Ezcurra. 2008. Do leaf margins of the temperate forest flora of southern South America reflect a warmer past? Global Ecology and Biogeography 17: 164–174.

Apostol, E. N., A. L. Curtu, L. M. Daia, B. Apostol, C. G. Dinu, and N. Şofletea. 2017. Leaf morphological variability and intraspecific taxonomic units for pedunculate oak and grayish oak (genus *Quercus* L., series *Pedunculatae* Schwz.) in Southern Carpathian Region (Romania). Science of The Total Environment 609: 497–505.

Auguie, B. 2017. gridExtra: Miscellaneous Functions for ‘Grid’ Graphics.

Bailey, I. W., and E. W. Sinnott. 1915. A Botanical Index of Cretaceous and Tertiary Climates. Science 41:831–834.

Bailey, I. W., and E. W. Sinnott. 1916. The Climatic Distribution of Certain Types of Angiosperm Leaves. American Journal of Botany 3: 24–39.

Baranski, M. J. 1975. An analysis of variation within white oak (*Quercus alba* L.). North Carolina Agricultural Experiment Station, Raleigh.

Becker, R. A., A. R. Wilks, R. Brownrigg, T. P. Minka, and A. Deckmyn. 2018. maps: Draw Geographical Maps.

Bivand, R. S., E. Pebesma, and V. Gomez-Rubio. 2013. Applied spatial data analysis with R, Second edition. Springer, NY.

Blue, M. P., and R. J. Jensen. 1988. Positional and seasonal variation in oak (*Quercus*, Fagaceae) leaf morphology. American Journal of Botany 75: 939–947.

Bonhomme, V., S. Picq, C. Gaucherel, and J. Claude. 2014. Momocs: Outline Analysis Using R. Journal of Statistical Software 56: 1–24.

Bragg, J. G., and M. Westoby. 2002. Leaf size and foraging for light in a sclerophyll woodland. Functional Ecology 16: 633–639.

Bruschi, P., P. Grossoni, and F. Bussotti. 2003. Within-and among-tree variation in leaf morphology of *Quercus petraea* (Matt.) Liebl. natural populations. Trees 17: 164–172.

Burnham, R. J., and G. S. Tonkovich. 2011. Climate, leaves, and the legacy of two giants. New Phytologist 190: 514–517.

Cavender-Bares, J. 2019. Diversification, adaptation, and community assembly of the American oaks (*Quercus*) a model clade for integrating ecology and evolution. New Phytologist 221: 669–692.

Crampton, J. S. 1995. Elliptic Fourier shape analysis of fossil bivalves: some practical considerations. Lethaia 28: 179–186.

Ducrey, M. 1992. Variation in leaf morphology and branching pattern of some tropical rain forest species from Guadeloupe (French West Indies) under semi-controlled light conditions. Annales des Sciences Forestières 49: 553–570.

García□Nogales, A., J. C. Linares, R. G. Laureano, J. I. Seco, and J. Merino. 2016. Range-wide variation in life-history phenotypes: spatiotemporal plasticity across the latitudinal gradient of the evergreen oak *Quercus ilex*. Journal of Biogeography 43: 2366–2379.

Garner, M., K. K. Pham, A. T. Whittemore, J. Cavender-Bares, P. F. Gugger, P. S. Manos, I. S. Pearse, and A. L. Hipp. 2019. From Manitoba to Texas: A study of the population genetic structure of bur oak (*Quercus macrocarpa*). International Oaks: The Journal of the International Oak Society 30: 131–138.

Genz, A., and F. Bretz. 2009. Computation of Multivariate Normal and t Probabilities. Springer-Verlag, Heidelberg.

Genz, A., F. Bretz, T. Miwa, X. Mi, F. Leisch, F. Scheipl, and T. Hothorn. 2019. mvtnorm: Multivariate Normal and t Distributions.

Givnish, T. J. 1987. Comparative Studies of Leaf Form: Assessing the Relative Roles of Selective Pressures and Phylogenetic Constraints. New Phytologist 106: 131–160.

González-Rodríguez, A., D. M. Arias, S. Valencia-Avalos, and K. Oyama. 2004. Morphological and RAPD analysis of hybridization between *Quercus affinis* and *Q. laurina* (Fagaceae), two Mexican red oaks. American Journal of Botany 91: 401–409.

González-Rodríguez, A., and K. Oyama. 2005. Leaf morphometric variation in *Quercus affinis* and *Q. laurina* (Fagaceae), two hybridizing Mexican red oaks. Botanical Journal of the Linnean Society 147:427–435.

Greenwood, D. R., P. Wilf, S. L. Wing, and D. C. Christophel. 2004. Paleotemperature Estimation Using Leaf-Margin Analysis: Is Australia Different? PALAIOS 19: 129–142.

Halloy, S. R. P., and A. F. Mark. 1996. Comparative leaf morphology spectra of plant communities in New Zealand, the Andes and the European Alps. Journal of the Royal Society of New Zealand 26: 41–78.

Hamerlynck, E. P., and A. K. Knapp. 1994. Leaf-level responses to light and temperature in two co-occurring *Quercus* (Fagaceae) species: implications for tree distribution patterns. Forest Ecology and Management 68: 149–159.

Haralick, R. M. 1974. A Measure for Circularity of Digital Figures. IEEE Transactions on Systems, Man, and Cybernetics SMC-4: 394–396.

Hijmans, R. 2017. raster: Geographic Data Analysis and Modeling. R package version 2.6-7.

Hipp, A. L., P. S. Manos, A. González-Rodríguez, M. Hahn, M. Kaproth, J. D. McVay, S. V. Avalos, and J. Cavender-Bares. 2018. Sympatric parallel diversification of major oak clades in the Americas and the origins of Mexican species diversity. New Phytologist 217: 439–452.

Hipp, A. L., A. T. Whittemore, M. Garner, M. Hahn, E. Fitzek, E. Guichoux, J. Cavender-Bares, et al. 2019. Genomic identity of white oak species in an eastern North American syngameon. Annals of the Missouri Botanical Garden 104: 455–477.

Hoban, S. 2014. An overview of the utility of population simulation software in molecular ecology. Molecular Ecology 23: 2383–2401.

Hoban, S. 2019. New guidance for ex situ gene conservation: Sampling realistic population systems and accounting for collection attrition. Biological Conservation 235: 199–208.

Hoban, S., T. Callicrate, J. Clark, S. Deans, M. Dosmann, J. Fant, O. Gailing, et al. 2020. Taxonomic similarity does not predict necessary sample size for ex situ conservation: a comparison among five genera. Proceedings of the Royal Society B: Biological Sciences 287: 20200102.

Hoban, S., O. Gaggiotti, and G. Bertorelle. 2013. Sample Planning Optimization Tool for conservation and population Genetics (SPOTG): a software for choosing the appropriate number of markers and samples. Methods in Ecology and Evolution 4: 299–303.

Howard, D., R. Preszler, J. Williams, S. Fenchel, and W. Boecklen. 1997. How discrete are oak species? Insights from a hybrid zone between *Quercus grisea* and *Quercus gambelii*. Evolution 51: 747–755.

Jensen, R. J. 1990. Detecting shape variation in oak leaf morphology: A comparison of rotational-fit methods. American Journal of Botany 77: 1279–1293.

Jensen, R. J., R. Depiero, and B. K. Smith. 1984. Vegetative characters, population variation, and the hybrid origin of *Quercus ellipsoidalis*. American Midland Naturalist 111: 364–370.

Kaproth, M. A., and J. Cavender-Bares. 2016. Drought tolerance and climatic distributions of the American oaks. International Oak Journal 27: 49–60.

Koenig, W. D., J. M. H. Knops, J. L. Dickinson, and B. Zuckerberg. 2009. Latitudinal decrease in acorn size in bur oak (*Quercus macrocarpa*) is due to environmental constraints, not avian dispersal. Botany 87: 349–356.

Kremer, A., J. Dupouey, J. Deans, J. Cottrell, U. Csaikl, R. Finkeldey, S. Espinel, et al. 2002. Leaf morphological differentiation between *Quercus robur* and *Quercus petraea* is stable across western European mixed oak stands. Annals of Forest Science 59: 777–787.

Li, Y., P. B. Reich, B. Schmid, N. Shrestha, X. Feng, T. Lyu, B. S. Maitner, et al. 2020. Leaf size of woody dicots predicts ecosystem primary productivity. Ecology Letters 23: 1003–1013.

Little, E. L. 1971. Atlas of United States trees, volume 1, Conifers and important hardwoods. U.S. Dept. of Agriculture, Forest Service, Washington, D.C.□:

Little, S. A., S. W. Kembel, and P. Wilf. 2010. Paleotemperature Proxies from Leaf Fossils Reinterpreted in Light of Evolutionary History. PLOS ONE 5: e15161.

Liu, M., Z. Wang, S. Li, X. Lü, X. Wang, and X. Han. 2017. Changes in specific leaf area of dominant plants in temperate grasslands along a 2500-km transect in northern China. Scientific Reports 7: 10780.

Marron, N., E. Dreyer, E. Boudouresque, D. Delay, J.-M. Petit, F. M. Delmotte, and F. Brignolas. 2003. Impact of successive drought and re-watering cycles on growth and specific leaf area of two Populus × canadensis (Moench) clones, ‘Dorskamp’ and ‘Luisa_Avanzo’. 23: 11.

McCarthy, D. M., and R. J. Mason-Gamer. 2019. Morphological Variation in North American *Tilia* and Its Value in Species Delineation. International Journal of Plant Sciences 181: 175–195.

McDonald, P. G., C. R. Fonseca, J. M. Overton, and M. Westoby. 2003. Leaf-size divergence along rainfall and soil-nutrient gradients: is the method of size reduction common among clades? Functional Ecology 17: 50–57.

McKee, M. L., D. L. Royer, and H. M. Poulos. 2019. Experimental evidence for species-dependent responses in leaf shape to temperature: Implications for paleoclimate inference. PLOS ONE 14: e0218884.

McKee, M., and D. L. Royer. 2017. How Does Temperature Impact Leaf Size and Shape in Four Woody Dicot Species? Testing the Assumptions of Leaf Physiognomy-Climate Models. AGU Fall Meeting Abstracts 11.

Minchin, P. R. 1987. An evaluation of the relative robustness of techniques for ecological ordination. Vegetatio 69: 89–107.

Moles, A. T., S. E. Perkins, S. W. Laffan, H. Flores□Moreno, M. Awasthy, M. L. Tindall, L. Sack, et al. 2014. Which is a better predictor of plant traits: temperature or precipitation? Journal of Vegetation Science 25: 1167–1180.

Mooney, H. A., and E. L. Dunn. 1970. Convergent Evolution of Mediterranean-Climate Evergreen Sclerophyll Shrubs. Evolution 24: 292–303.

Oksanen, J., F. G. Blanchet, M. Friendly, R. Kindt, P. Legendre, D. McGlinn, P. R. Minchin, et al. 2017. vegan: Community Ecology Package. R package version 2.4-5.

Pebesma, E. J., and R. S. Bivand. 2005. Classes and methods for spatial data in R. R News 5: 9–13.

Peppe, D. J., D. L. Royer, B. Cariglino, S. Y. Oliver, S. Newman, E. Leight, G. Enikolopov, et al. 2011. Sensitivity of leaf size and shape to climate: global patterns and paleoclimatic applications. New Phytologist 190: 724–739.

R Core Team. 2018. R: A language and environment for statistical computing, version 3.4.4. R Foundation for Statistical Computing, Vienna.

Ramírez-Valiente, J. A., and J. Cavender-Bares. 2017. Evolutionary trade-offs between drought resistance mechanisms across a precipitation gradient in a seasonally dry tropical oak (*Quercus oleoides*). Tree Physiology: 1–13.

Ramírez-Valiente, J. A., A. Center, J. Sparks, K. Sparks, J. Etterson, T. Longwell, G. Pilz, and J. Cavender-Bares. 2017. Population-level differentiation in growth rates and leaf traits in seedlings of the neotropical live oak *Quercus oleoides* grown under natural and manipulated precipitation regimes. Frontiers in Plant Science 8.

Ramírez□Valiente, J. A., R. López, A. L. Hipp, and I. Aranda. 2020. Correlated evolution of morphology, gas exchange, growth rates and hydraulics as a response to precipitation and temperature regimes in oaks (*Quercus*). New Phytologist 227: 794–809.

Rodríguez-Correa, H., K. Oyama, I. MacGregor-Fors, and A. González-Rodríguez. 2015. How are oaks distributed in the neotropics? A perspective from species turnover, areas of endemism, and climatic niches. International Journal of Plant Sciences 176: 222–231.

Rosin, P. L. 2005. Computing global shape measures. Handbook of Pattern Recognition and Computer Vision, 177–196. WORLD SCIENTIFIC.

Royer, D. L., J. C. McElwain, J. M. Adams, and P. Wilf. 2008. Sensitivity of leaf size and shape to climate within *Acer rubrum* and *Quercus kelloggii*. New Phytologist 179: 808–817.

Royer, D. L., and P. Wilf. 2006. Why Do Toothed Leaves Correlate with Cold Climates? Gas Exchange at Leaf Margins Provides New Insights into a Classic Paleotemperature Proxy. International Journal of Plant Sciences 167: 11–18.

Royer, D. L., P. Wilf, D. A. Janesko, E. A. Kowalski, and D. L. Dilcher. 2005. Correlations of climate and plant ecology to leaf size and shape: potential proxies for the fossil record. American Journal of Botany 92: 1141–1151.

Schmerler, S. B., W. L. Clement, J. M. Beaulieu, D. S. Chatelet, L. Sack, M. J. Donoghue, and E. J. Edwards. 2012. Evolution of leaf form correlates with tropical-temperate transitions in Viburnum (Adoxaceae). Proceedings of the Royal Society B: Biological Sciences 279: 3905–3913.

Schnabel, A., and J. L. Hamrick. 1990. Comparative Analysis of Population Genetic Structure in Quercus macrocarpa and *Q. gambelii* (Fagaceae). Systematic Botany 15: 240–251.

Schneider, C. A., W. S. Rasband, and K. W. Eliceiri. 2012. NIH Image to ImageJ: 25 years of image analysis. Nature Methods 9: 671–675.

Sokal, R. R., T. J. Crovello, and R. S. Unnasch. 1986. Geographic Variation of Vegetative Characters of Populus deltoides. Systematic Botany 11: 419–432.

Stein, J., D. Binion, and R. Acciavatti. 2003. Field Guide to Native Oak Species of Eastern North America (FHTET-2003-01). United States Department of Agriculture Forest Service, Forest Health Technology Enterprise Team, Morgantown.

Tracey, S. R., J. M. Lyle, and G. Duhamel. 2006. Application of elliptical Fourier analysis of otolith form as a tool for stock identification. Fisheries Research 77: 138–147.

Walls, R. L. 2011. Angiosperm leaf vein patterns are linked to leaf functions in a global-scale data set. American Journal of Botany 98: 244–253.

Wickham, H. 2009. ggplot2: Elegant Graphics for Data Analysis. Springer-Verlag, New York.

Wright, I. J., N. Dong, V. Maire, I. C. Prentice, M. Westoby, S. Díaz, R. V. Gallagher, et al. 2017. Global climatic drivers of leaf size. Science 357: 917–921.

Wright, I. J., P. B. Reich, M. Westoby, D. D. Ackerly, Z. Baruch, F. Bongers, J. Cavender-Bares, et al. 2004. The worldwide leaf economics spectrum. Nature 428: 821–827.

