## Supplementary figures and images for "Leaf shape and size variation in bur oaks: An empirical study and simulation of sampling strategies"

### Appendix 1

Desmond et al. - American Journal of Botany 2021 - Appendix S1

Biplots of all simple regressions

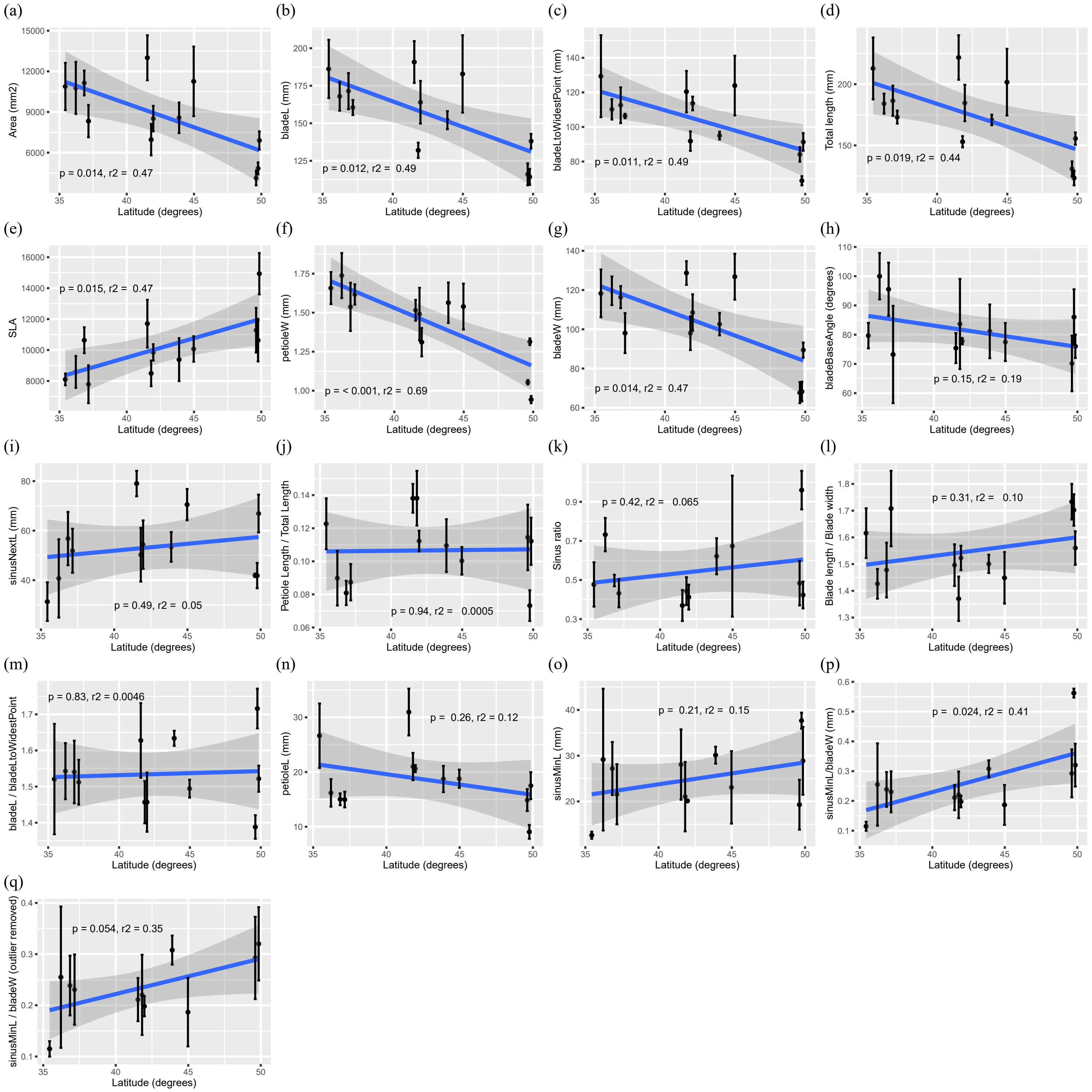
