## Appendix 2 for "Leaf shape and size variation in bur oaks: An empirical study and simulation of sampling strategies"

Desmond et al. - American Journal of Botany 2021 - Appendix S2
PCA based on eFourier analysis of leaf outlines

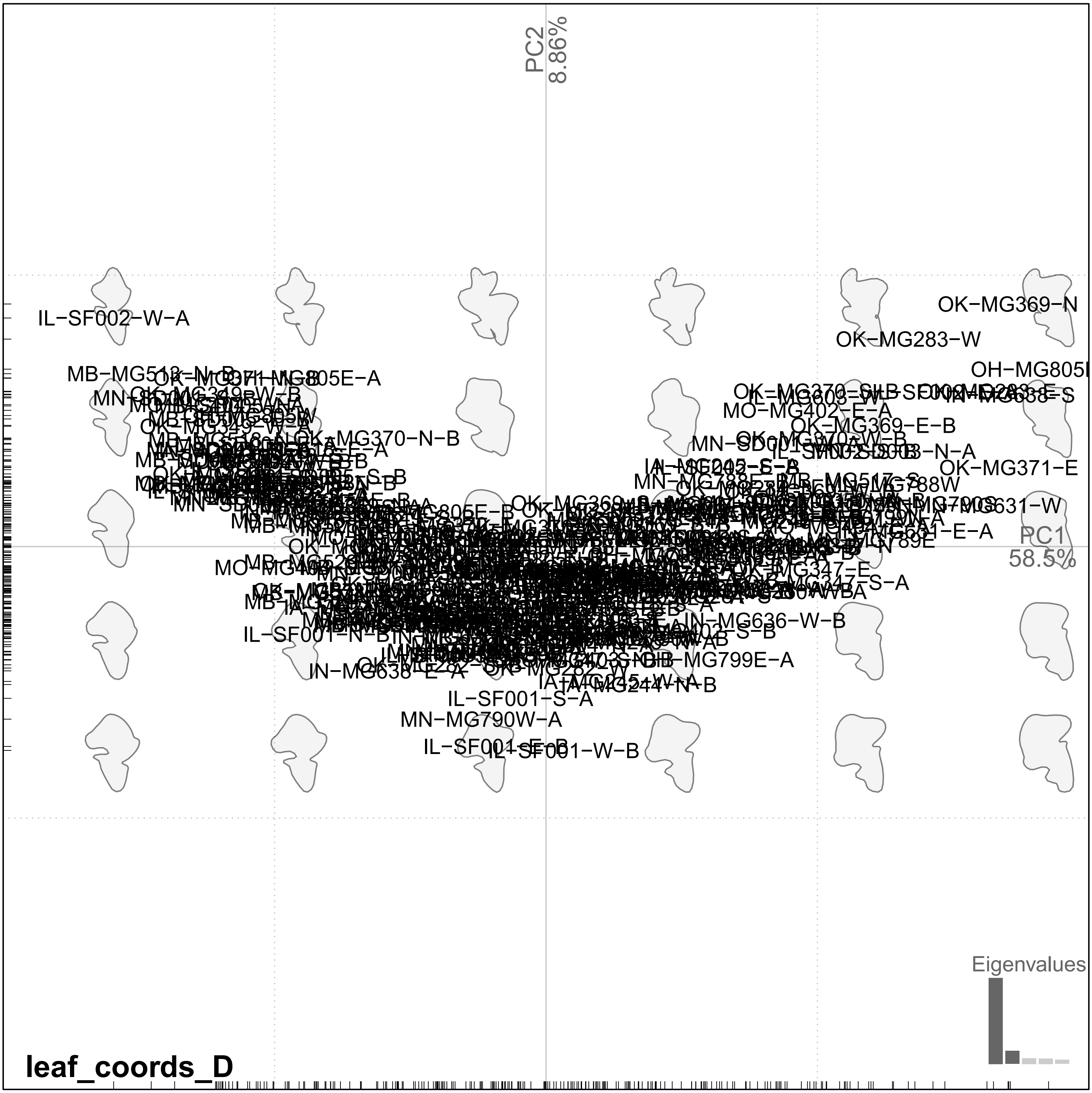
