## Appendix 3 for "Leaf shape and size variation in bur oaks: An empirical study and simulation of sampling strategies"

Desmond et al. - American Journal of Botany 2021 - Appendix S3
Non-metric multidimensional scaling ordination of leaf measurements

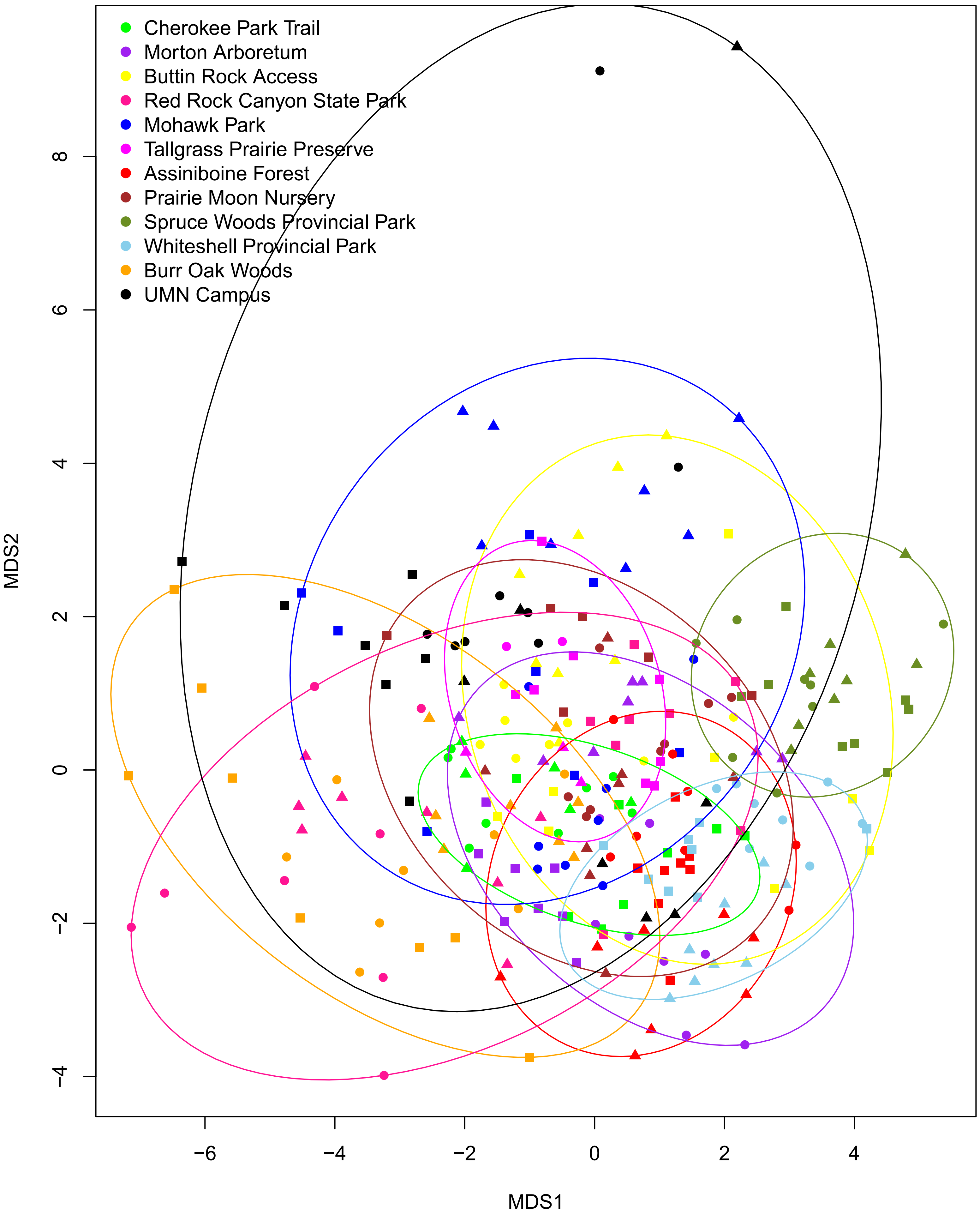
